# Microbiota-pathogen interactions after host death: a potential determinant of pathogen evolution

**DOI:** 10.64898/2026.08.25.746672

**Authors:** Alexandra von Bismarck, Haicheng Xie, Mathias Franz, Maryam Keshavarz

## Abstract

During the lifetime of many animals, the microbiota fulfills multiple functions. After death of their hosts, these microbes contribute to cadaver decomposition, with implications for forensics, fossilization and soil nutrient and microbial community dynamics. Here, we draw attention to the possibility that host microbiota can also influence the evolution of pathogen lifestyles. We hypothesized that competition between microbiota and pathogens after host death can reduce benefits to pathogens of killing and decomposing their host. To test this hypothesis, we conducted infection experiments in which we injected the entomopathogenic bacteria *Pseudomonas entomophila* into *Tenebrio molitor* larvae. Our results show that bacterial proliferation after pathogen-induced host death occurs in larvae with strongly reduced microbiota, but not in larvae with intact gut microbiota. Strikingly, we found that gut microbiota can suppress the proliferation of an about 100 times larger pathogen population. In addition, we identified a microbiota member that might have mediated competitive suppression of pathogen proliferation after host death. Taken together, our results support our hypothesis that decomposing host microbiota can effectively compete with pathogens, thereby reducing the fitness of pathogens that kill and then exploit dead hosts. Based on a reanalysis of an existing theoretical model, we conclude that the host microbiota can facilitate the evolution of more benign pathogens that are less likely to kill their host for cadaver exploitation. Thus, our findings highlight the potentially important but so far unexplored possibility that pathogen-microbiota interactions in dead hosts can affect living hosts by influencing the evolution of pathogens lifestyles.

**Significance Statement:** A variety of microbes colonize animals, especially their guts. Throughout the host’s lifetime, these microbes perform important functions, including defense against pathogens. Following host death, some of these members also contribute to cadaver decomposition. Here, we propose that the role of host-associated microbes in host decomposition could influence pathogen evolution. In experimental infections of mealworm larvae with a bacterial pathogen, we found that gut microbes limit pathogen proliferation in mealworm cadavers. Combined with a mathematical model, these findings reveal a previously unrecognized influence on pathogen evolution: microbial competition after host death reduces the benefits of killing the host, potentially favoring the evolution of more benign pathogens.

## Introduction

The life of animals can be profoundly impacted by their microbiota via influences on nutrition (Hacquard et al., 2015; Jang et al., 2025), growth and development (Perlman et al., 2022), behavior (Liberti & Engel, 2020; Vuong et al., 2017) and protection against pathogens (Chiu et al., 2017; Kamada et al., 2013). After host death these microbes, especially from the gut, contribute to cadaver decomposition (Martino et al., 2022; Preiswerk et al., 2018), with implications for soil nutrient and microbial dynamics (Carter et al., 2007; DeBruyn et al., 2025), fossilization (Butler et al., 2015) and forensics (Metcalf et al., 2013; Metcalf et al., 2016). Here we want to draw attention to another potential implication of host-microbe associations after host death: the evolution of pathogen lifestyles.

The spectrum of pathogenic lifestyles ranges from mildly or moderately virulent species that transmit from living hosts (Kennedy, 2023; Walther & Ewald, 2004) to highly virulent obligate killers that require host death for propagation (Ebert et al., 1996; Evans, 1982; Miller et al., 1983). Along this spectrum, virulence can be generally defined as reduction in host fitness due to the pathogen infection (Read, 1994). Theoretical studies, in which virulence is usually conceived as the added host death rate during an infection (Alizon et al., 2009; Anderson & May, 1982; Cressler et al., 2016; Day, 2002; Frank, 1996), have focused on assessing how virulence evolves for a given pathogen lifestyle, including direct transmission from living hosts (Alizon & Lion, 2011; Alizon & van Baalen, 2005; Anderson & May, 1982; Ganusov et al., 2002; Gilchrist & Coombs, 2006), transmission from dead host in obligate killers (Ebert & Weisser, 1997; MacDonald & Brisson, 2023; Redman et al., 2016) or mixed transmission from dead and living hosts (Day, 2002). Empirical studies in this context have focused on testing the general prediction that trade-offs between virulence and transmission shape the evolution of virulence (e.g. Acevedo et al., 2019). However, what determines evolutionary transitions among different pathogen lifestyles remains poorly understood. Here, we propose that transitions between pathogens transmitting from living hosts and pathogens killing their hosts to decompose and transmit from dead hosts can be influenced by the host gut microbiota.

Our main hypothesis is that some members of gut microbiota can reduce benefits for the pathogen of killing and decomposing its host due to competition over decomposition of the dead host. To test this hypothesis, we conducted experiments on larvae of the mealworm beetles *Tenebrio molitor* and the entomopathogen *Pseudomonas entomophila*. *P. entomophila* was first isolated from *Drosophila melanogaster* (Vodovar et al., 2005) and is highly pathogenic to this host upon both ingestion and injection (Dieppois et al., 2014; Hidalgo et al., 2022). Following experimental injection, this bacterium can also infect and kill insects from other orders, including *T. molitor* and *Galleria mellonella* (Dieppois et al., 2014; Keshavarz et al., 2023; Kordaczuk et al., 2025), making it a key model for studying insect–pathogen interactions (Dieppois et al., 2014; Henry et al., 2025). To assess potential evolutionary implications of our findings we additionally reanalysed an existing mathematical model of pathogen virulence evolution (Day, 2002).

## Results

### Microbiota contributes to cadaver decomposing in *T. molitor* larvae

We first confirmed that the host microbiota contributes to cadaver decomposition in *T. molitor* larvae that were killed via snap freezing. To account for substantial variation in recorded microbiota load, we fitted a zero-inflated generalized linear mixed model. This model indicated that microbiota load increased over time in dead larvae (Fig. 1A). Specifically, microbiota load among non-zero load values increased more than tenfold within three days (*p*<0.001, *z*=8.070), whereas we did not find a statistically significant effect for the proportion of zeros loads (*p*=0.315, *z*=-1.005).

**Fig. 1:**
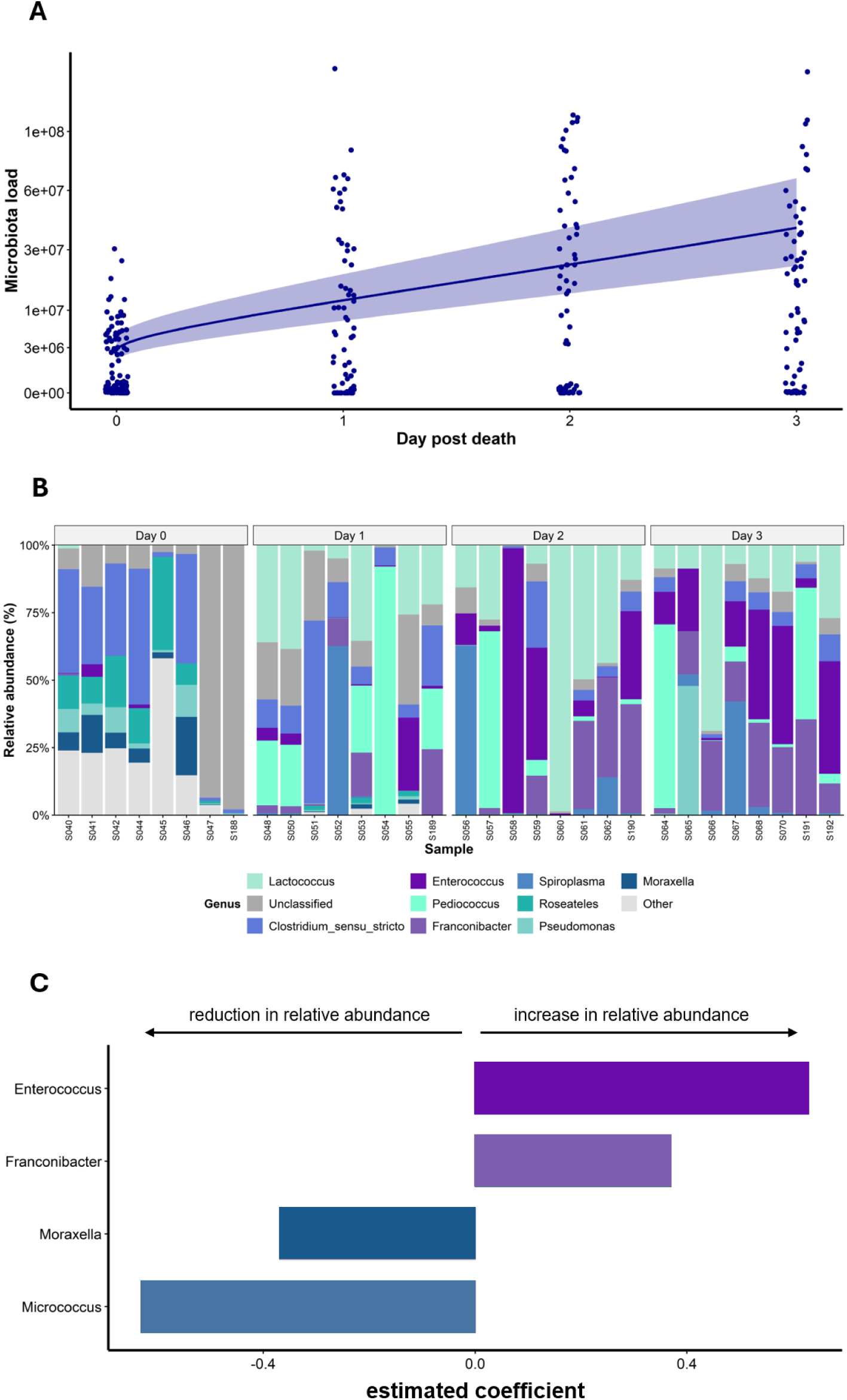
(**A**) Microbiota load after artificial death of *T. molitor* larvae by snap freezing. Each dot represents the measured microbiota load of a single larva from four independent replicates. Points are jittered on the x-axis to increase visibility. The line and shaded area show model estimates for temporal dynamics with the corresponding 95% confidence interval. A box-Cox transformation was applied to the y-axis to appropriately capture the variation among data points (Box & Cox, 1964). (**B**) Genus-level relative community composition of the 10 most abundant taxa in dead *T. molitor* larvae, determined by 16S rRNA gene sequencing. Each bar represents an individual larva; colored segments denote distinct taxa, with segment height being proportional to relative abundance within that sample. (**C**) Results of compositional data analysis (Calle et al., 2023) identified two microbiota taxa that increased in relative abundance after host death (indicated by positive coefficients) and two taxa that decreased in relative abundance during 3 days after host death (indicated by with negative coefficients). Absolute values of coefficients indicate how the magnitude of change in each taxon relates to the magnitude of changes in other selected taxa.

We applied compositional data analysis to detect temporal shifts in gut microbial composition during three days after host death (Fig. 1B), identifying four taxa whose relative abundance changed over time. Specifically, relative abundances of *Enterococcus* and *Franconibacter* increased, whereas *Moraxella* and *Micrococcus* decreased (Fig. 1B, C).

### Microbiota limits *P. entomophila* proliferation after pathogen induced host death

To test our hypothesis regarding potential interactions between host gut microbiota and the pathogen after pathogen-induced host death, we systemically infected *T. molitor* larvae fed on different diets: antibiotic-treated (AB) larvae, which strongly suppresses host gut microbiota (Keshavarz et al., 2025), and untreated larvae as controls (CR). Following natural death after infection, the recorded pathogen load showed substantial variation, with temporal dynamics depending on dietary treatment (Fig. 2A). On the day of death, the pathogen load was about two orders of magnitudes larger than the recorded microbiota load after snap freezing (Fig. 1A) or living larvae (Fig. S1). In addition, the pathogen load was on average about 1.6 times higher in AB treatment compared to CR (result for diet: *p*=0.021, *z*=-2.310). The further temporal dynamics after host death differed between dietary treatments as indicated by a significant interaction between diet and day (*p*<0.001, *z*=-3.730). Specifically, estimated changes in pathogen load within three days were 1) an increase of about 5.3-foldin the AB treatment, and 2) a decrease to approximately 85% in the CR treatment. Taken together, these results support our hypothesis that the host gut microbiota limits pathogen proliferation after host death.

**Fig. 2:**
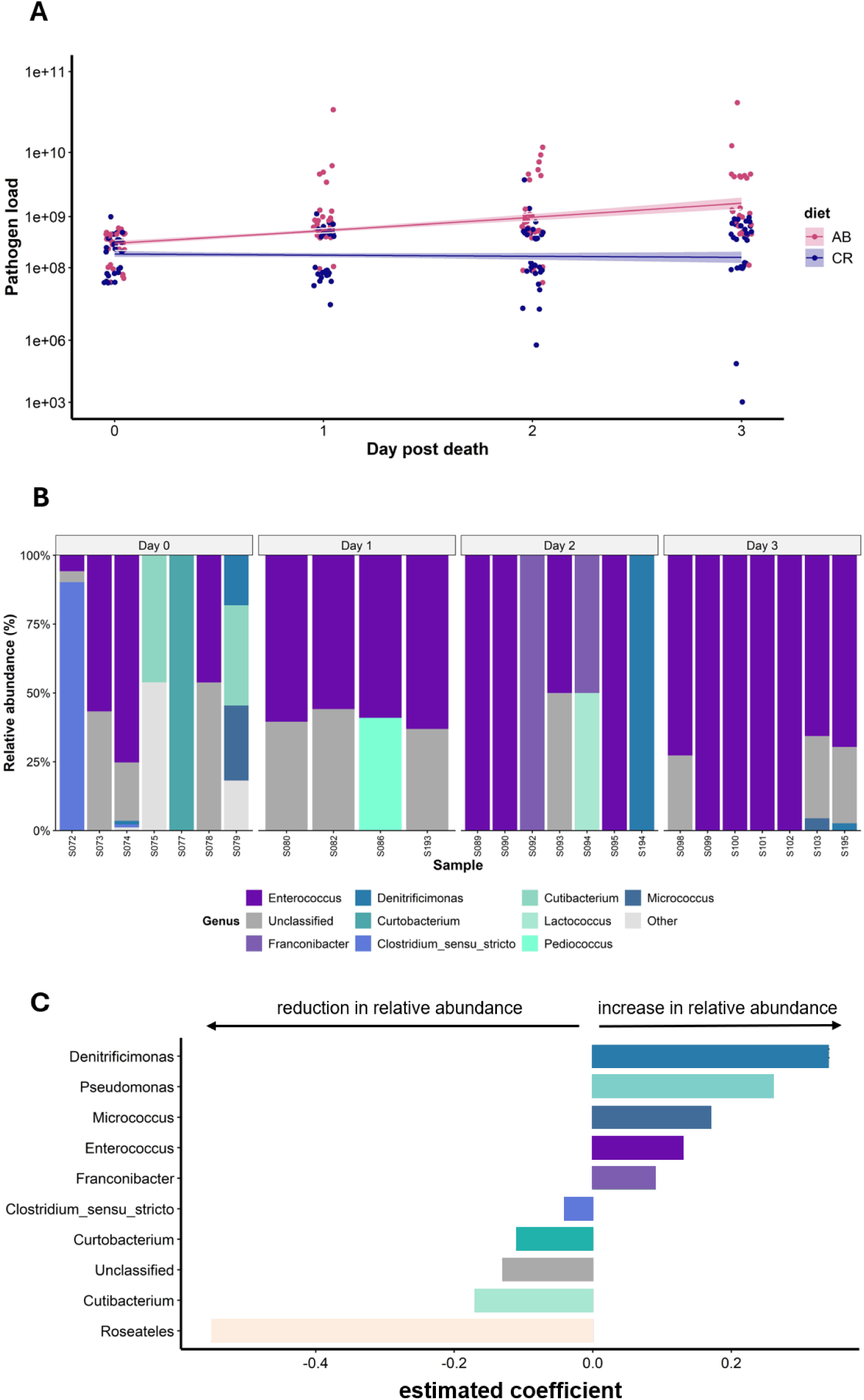
(**A**) Pathogen load after natural death of *T. molitor* larvae infected with *P. entomophila* under two treatments that modulate the host gut microbiota: antibiotic-treated (AB) larvae, which strongly suppresses host gut microbiota (Keshavarz et al., 2025), and conventional larvae as controls (CR). Each dot represents the pathogen load of a single larva from three independent replicates. Points are jittered on the x-axis to increase visibility. Lines and shaded areas show model estimates for temporal dynamics with the corresponding 95% confidence intervals. A box-Cox transformation was applied to the y-axis to appropriately capture the variation among data points (Box & Cox, 1964). (**B**) Genus-level relative community composition of 10 dominant taxa in *P. entomophila*-infected larvae, after removing *P. entomophila*, determined by 16S rRNA gene sequencing. Each bar represents an individual larva; colored segments denote distinct taxa, with segment height being proportional to relative abundance within that sample. (**C**) Results of compositional data analysis (Calle et al., 2023) identified microbiota taxa that increased or decreased in relative abundance during three days after host death. Absolute values of coefficients indicate how the magnitude of change in each taxon relates to the magnitude of changes in other selected taxa

In addition, compositional data analysis of microbiota composition during 3 days after host death (Fig. 2B) identified 10 taxa whose relative abundance changed over time. Among these, relative abundance of *Denitrificimonas*, *Pseudomonas*, *Micrococcus*, *Enterococcus*, and *Franconibacter* increased over time, whereas relative abundance of *Clostridium*_*sensu*_*stricto*, *Curtobacterium*, *Cutibacterium*, *Roseateles* decreased over time (Fig. 2C).

### Enterococcus sp. inhibits P. entomophila proliferation in vitro

Using a cultured isolate of *Enterococcus* sp. from the *T. molitor* gut, we tested its interspecific interactions with *P. e in vitro* in a modified cross-streak assay. In this qualitative assay, gut-derived microbiota was streaked perpendicularly, and *P. e* was then introduced into the adjacent quadrants to assess inhibition. Our results revealed that *Enterococcus* sp. inhibited the growth of *P. e* (Fig. 3, Fig. S2).

**Fig. 3:**
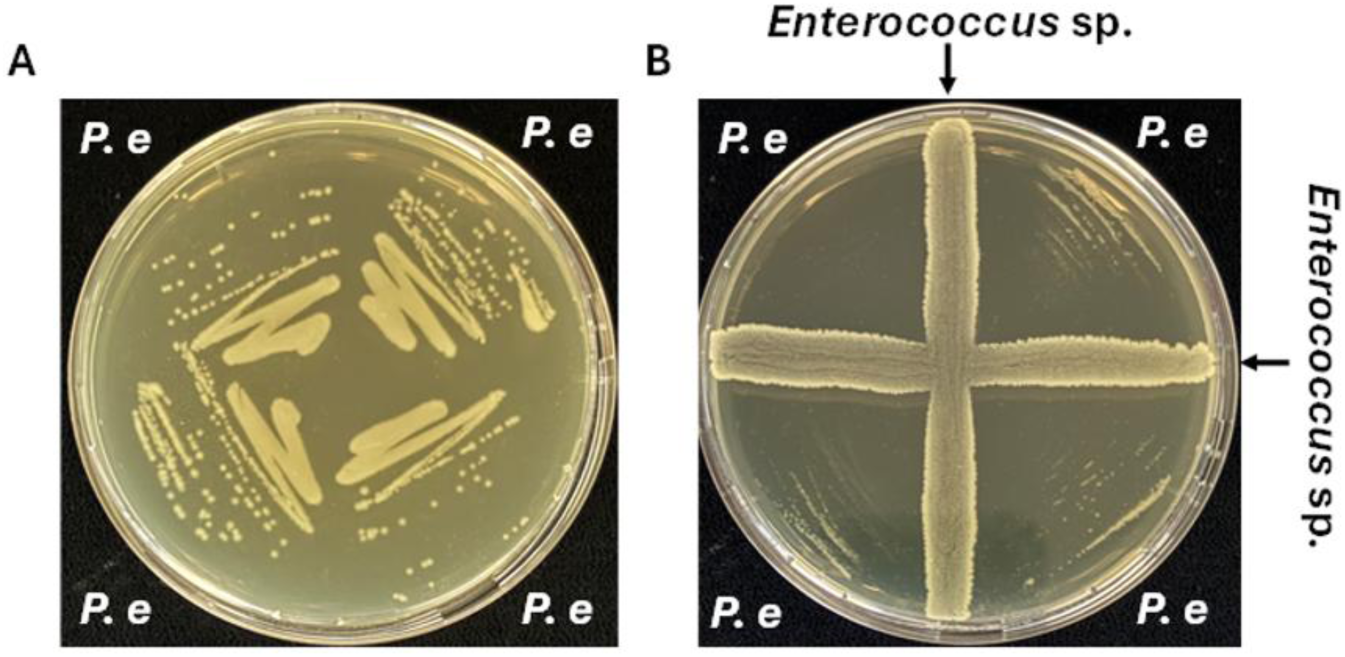
Modified cross-streak assay of gut-derived *Enterococcus* sp. against *P. e*. (**A**) *P. e* grown in monoculture (control). (**B**) Inhibition of *P. e* growth by *Enterococcus* sp., indicated by a zone of inhibition adjacent to the *Enterococcus* streak in binary culture. Results of 11 additional plates from two independent experiments are included in the Supporting Information (Fig. S2).

### Theoretical analysis

To test potential evolutionary implications of our empirical finding that the host gut microbiota inhibits pathogen proliferation after host death, we reanalyzed the mathematical virulence evolution model of Day (2002). Although this model was used to explore the consequences of spore and toxin production for the evolution of pathogen-induced host mortality, the model is also suitable for investigating the expected effects of microbiota-induced reduction in pathogen proliferation after host death (see Methods for more details).

## Discussion

In this study, we experimentally investigated the interaction between the entomopathogen *P. entomophila* and the gut microbiota of *T. molitor* larvae after host death. After establishing that the gut microbiota contributes to cadaver decomposition in uninfected hosts (Fig. 1), our investigation in infected hosts revealed that pathogen proliferation after host death is constrained by the gut microbiota (Fig. 2). This result aligns with our prediction and thus supports our hypothesis that the host gut microbiota can compete with pathogens that kill their host, thereby limiting the benefits to the pathogen to kill and decompose its hosts. To our knowledge, this finding emphasizes an unexplored effect of the host gut microbiota on the evolution of pathogen lifestyles.

It is already well recognized that pathogen evolution can be influenced by different forms of competition between pathogens and microbiota in living hosts (Armitage et al., 2022). Based on our results, we propose that such an influence of the host microbiota can also occur after host death. Specifically, we suggest that competition of the host gut microbiota with pathogens in dead hosts could exert selection pressures on pathogens that oppose the evolution of host killing for the purpose of cadaver decomposition. This effect is likely to be particularly relevant for pathogens that are able to transmit from living as well as dead hosts, such as the entomopathogenic fungus *Beauveria bassiana*, the bacterium *Bacillus thuringiensis* (Milutinović et al., 2015) and bacterial pathogens in fish farms (Pulkkinen et al., 2010). In such cases, if the host microbiota competes with the pathogen after host death, this should decrease selection for host killing. Alternatively, it could select for later killing that keeps the host alive longer and thus avoid or delay competition with the host microbiota. This argument is also supported by an initial theoretical investigation using the mathematical model of Day (2002) on pathogen virulence evolution (Fig. 4). Accordingly, competition between host microbiota and pathogens after host death should act against a pathogen lifestyle that involves killing and exploiting dead hosts. Future theoretical work could explore this idea in greater depth. In addition, empirical studies could test this idea more directly by investigating pathogens that transmit both from living and dead hosts.

**Fig. 4:**
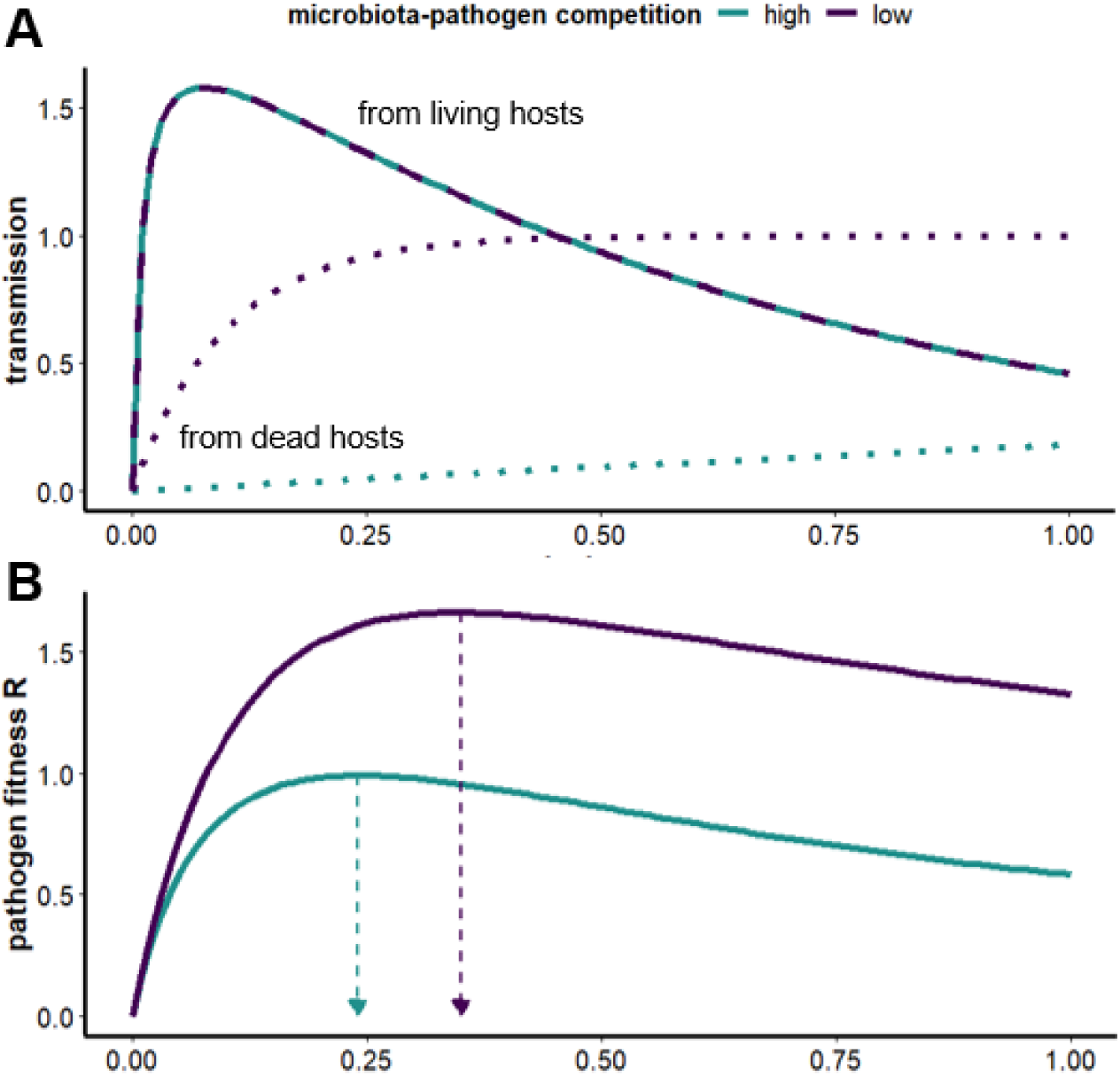
Predictions from a reanalyzed mathematical model of Day (2002) on the influence of competition between microbiota and pathogen after host death. (**A)** Model assumptions: pathogens can transmit from living (dashed lines) and dead hosts (dotted lines), and we additionally assumed that competition with the microbiota reduces transmission from dead hosts (green dotted line compared to purple dotted line). (**B)** Effect on pathogen fitness: solid lines depict pathogen fitness along a virulence gradient, and corresponding dotted arrows indicate virulence levels that maximize fitness. See Methods for model details and chosen parameter values.

Our results provide some initial insights in to how the host microbiota competes with *P. entomophila* in dead *T. molitor* larvae. The host microbiota inhibited pathogen growth after host death even though its recorded load was approximately 100-fold lower than the pathogen load (Fig. 1, 2), which indicates a high efficacy of the underlying competitive effect. Our analysis of changes in microbiota composition after host death identified two potential taxa that could have mediated the observed competitive effects: *Enterococcus* and *Franconibacter*. The relative abundance of both taxa increased in snap-frozen and *P. entomophila*-infected larvae, suggesting a potential role in host decomposition and pathogen growth inhibition (Fig. 1C, 2C). *Enterococcus*, the more abundant taxa, has been previously reported to be part of the gut microbiota in living *Tenebrio* (Bento de Carvalho et al., 2026; Osimani et al., 2018). Our modified cross-streak assay confirmed that *Enterococcus* sp. inhibits *P. entomophila in vitro* (Fig. 3), which due to the spatially distant effect most likely indicate interference competition. A potential mechanism for such a competitive effect could be environmental acidification via lactic acid production, which has been shown to be a mechanism how *Enterococcus faecalis* inhibit growth of *Pseudomoans aeruginosa* (Tan et al., 2022). Future studies are required to investigate the nature of the competitive mechanisms in more detail and confirm their occurrence *in vivo*.

Negative effects of the microbiota on pathogen growth dynamics after death might depend on pathogen infection route. In our experiments we injected the pathogen into the hemocoel to avoid potentially confounding effects of microbiota-pathogen interactions before death. Of note, injection-like exposure to pathogens can occur naturally through wounding can be common in the wild (Subasi et al., 2024), for example via ectoparasitic mites or traumatic insemination (Reinhardt et al., 2015; Stone et al., 2024). However, this setup might also have allowed the pathogen to reach very large numbers by the time of host death. Other infection routes, such as oral infections, might allow microbiota to compete with the pathogen at earlier stages, for example, while both occupy the gut lumen. Such competitive effects could limit pathogen load at death, further reducing the benefits of killing and exploiting dead hosts.

Not all microbiota-host associations are beneficial; they can range from commensal or mutualistic to pathogenic, with different implications for the evolution of pathogens. The ‘microbiome mutiny hypothesis’ proposes that in old or seriously ill people, some members of the human microbiota can switch from a benign, mutualistic lifestyle towards a highly virulent, saprophytic lifestyle that benefits form host death and its decomposition (Rózsa et al., 2015; Rózsa et al., 2017). Similarly, Preiswerk et al. (2018) suggested that microbiota members that are able to switch to a decomposing lifestyle after host death might facilitate the evolution of pathogens that exploit dead hosts. However, the opposite effects might also occur as suggested by our results: the ability of microbiota members to effectively compete with pathogens after host death could act against the evolution of pathogens that kill and exploit dead hosts. Further work is required to assess which of these possibilities is more likely and which factors affect the relative strength of these opposing effects.

## Methods

### *Tenebrio* husbandry

Late instar larvae of *Tenebrio molitor* (18^th^ to 20^th^ instar, approximately 2.5 - 3 cm in length) were acquired from a commercial supplier (Reptile Food Handels-u. Zucht GmbH, Berlin, Germany) (Park et al., 2014). Larvae were maintained at a population density of 500 larvae per container in the dark at 25 ± 3°C and 60±5% relative humidity (RH), provided with an *ad libitum* supply composed of autoclaved wheat bran, serving as their primary food source, along with a fresh apple slice every 2 days for hydration and supplementary nutrition. At each feeding, pupae were collected from the colony and kept in separate containers. Newly emerged adults were then transferred to separate containers for oviposition.

### Larval rearing diets and antibiotic treatment

Conventional (CR) early instar larvae (3^rd^ to 4^th^ instar, approximately 0.5 cm in length) were reared on a standardized basal diet consisting of 100 g of wheat bran, 10 g soy protein, 20 g soy flour, 170 g of a 5% yeast-wheat flour mixture, hydrated with 200 mL distilled water containing 0.2 mL propionic acid, a formulation used across all treatments to ensure nutritional consistency. For antibiotic-treated (AB) larvae, the same formulated diet was supplemented with 0.5 g of chloramphenicol, 0.5 g of sorbic acid, 0.5 mL of propionic acid in 200 mL of distilled water for 13 days. To mitigate the potential effects of antibiotic, larvae were subsequently fed on an identical diet devoid of chloramphenicol for an additional 2 days. The effectiveness of chloramphenicol on gut microbiota composition was previously performed using both culture-dependent and culture-independent approaches (Keshavarz et al., 2025). Fresh food was provided every 2 days to avoid desiccation.

### Bacterial culture

The following bacterial species were used in this study. *Pseudomonas entomophila* L48 (*P. e*) was retrieved from a 50% glycerol stock and cultivated in Luria-Bertani (LB) medium under aerobic conditions at 30°C for 16 hours with agitation. Optical density at 600 nm (OD600) was measured to obtain bacterial concentrations of 5 × 10⁷ colony forming units (CFU/mL), corresponding to an approximate injection dose of 10⁵ CFU per larva.

*Enterococcus* sp. was isolated from 10^th^ to 12^th^ instar larvae. Briefly, homogenates from individual larvae were spread onto Trypticase soy agar (TSA) plates. A single colony was re-streaked to obtain pure culture and identified by sequencing a partial region of the 16S rRNA gene using universal primers (27F/1492R). High-quality amplicon sequence variants (ASVs) were resolved using the DADA2 pipeline, and the resulting partial 16S rRNA gene sequences were assigned to taxa via BLASTn search against the NCBI nucleotide database (https://blast.ncbi.nlm.nih.gov/Blast.cgi?PAGE_TYPE=BlastSearch&BLAST_SPEC=MicrobialGe nomes). The partial 16S rRNA gene sequence is provided in the supplemental material. *Enterococcus* sp. was cultivated in LB medium under aerobic conditions at 37°C for 16 hours with agitation.

### Bacterial load

For quantifying microbial load, dead Individuals were placed in separate wells of square 100 mm Petri dishes (Sterilin™, Thermo Fisher Scientific), sealed with Parafilm, and maintained in a rearing incubator (at 25 ± 3°C and 60±5% RH), until plating. Each larva was homogenised in 100 µL LB broth using two sterile steel beads at 30 Hz for 20 seconds using a tissue homogenizer (Mill MM400, Retsch). Homogenates were serially diluted (1:10 to 1:10^5^) and plated onto the Trypticase soy agar (TSA) plates and incubated at 37°C for 16 hours. For bacterial load, the same procedure was followed, except that isolated *P. entomophila* were plated onto LB medium containing 1000 µg/mL ampicillin at 30°C for 16 hours (Keshavarz et al., 2023). Samples with no CFU at the lowest dilution were recorded as zero.

### DNA extraction and 16S rRNA gene amplicon sequencing

DNA isolation from each sample (from both alive and dead larvae) was performed using a modified protocol of the DNeasy PowerSoil Pro Kit (Qiagen, Germany), as previously described (Keshavarz et al., 2025). Briefly, homogenates in 800 µL of CD1 solution buffer were centrifuged at 25 Hz for 10 minutes, and incubated overnight with 10 μl of Proteinase K at 56°C for 16 hours in a ThermoMixer^®^ C. The remaining steps were carried out according to the manufacturer’s protocol. DNA concentration was measured via Qubit™ fluorometer (Invitrogen) and adjusted to 10 ng/µL. Amplification of the V4 region of the 16S rRNA gene was performed using primers 515F (5′-GTGYCAGCMGCCGCGGTA-3′) and 806R (5′-GGACTACNVGGGTWTCTAAT-3′). Libraries were prepared using a dual-index PCR approach with Q5 High-Fidelity DNA Polymerase (Keshavarz et al., 2025). Sequencing was carried out on an Illumina MiSeq platform using a 600-cycle v3 kit at the Berlin Center for Genomics in Biodiversity Research (BeGenDiv).

All bioinformatic processing was performed in software R (Team, 2020). Primers were removed from raw reads using cutadapt (Martin, 2011). Quality assessment of the primer-trimmed reads was performed with FastQC (Andrews et al., 2010). Based on the visualized quality profiles, reads were truncated to 200 bp for forward and 150 bp for reverse. Filtering and trimming were executed using the filterAndTrim function of the ‘dada2’ package (Callahan et al., 2016). Divisive Amplicon Denoising Algorithm 2 (DADA2) was then used to learn error rates, infer amplicon sequence variants (ASVs), and merge paired end reads. Taxonomy was assigned via DADA2’s naive Bayesian classifier (Wang et al., 2007) against the Ribosomal Database Project (RDP) training set v19 (Cole et al., 2014; Wang & Cole, 2024). ASVs classified as Eukaryota, Chloroplasts, or Mitochondria were excluded from further analyses (Hanshew et al., 2013).

### Experimental setup

#### Gut microbiota abundance and composition without pathogens

To assess whether culturable gut microbiota populations increase after death, 10^th^ to 12^th^ instar conventional larvae (approximately 1.3 - 1.8 cm in length) were snap-frozen in liquid nitrogen and sampled at 0, 1, 2, and 3 days post-death. Day 0 larvae were plated immediately after snap-freezing; larvae for days 1, 2, and 3 were held under standard rearing conditions until plating. Live larvae homogenized without freezing serving as controls for snap-freezing. We found no indication that snap freezing reduced microbiota load compared to living larvae (Fig. S1). Sixteen individual larvae per time point were processed in each of four independent experiments. Further, to determine the gut microbiota composition via 16S rRNA gene sequencing, 8 individual larvae per time point were bead-beaten for DNA extraction.

#### *Pseudomonas entomophila* proliferation and gut microbiota composition after pathogen induced host death

To test whether resource competition from the gut microbiota affects pathogen proliferation after death, CR- and AB-treated larvae (10^th^ to 12^th^ instar) were infected with 2 µL of *P. entomophila* (∼10⁵ CFU per larva). At 24 h post-infection, dead larvae were collected at 0 (i.e., at the time of death), 1, 2, and 3 days post-death. Day 0 larvae were plated immediately upon collection; larvae for later time points maintained under standard rearing conditions until plating. Ten larvae per time point were processed in each of three independent experiments. Next, for 16S rRNA gene sequencing, an additional 8 larvae per time point per treatment were homogenized by bead beating for DNA extraction.

### Microbe-microbe interaction

To test whether *Enterococcus* sp. inhibit *P. entomophila* growth *in vitro, Enterococcus* sp. was first streaked perpendicularly the center of TSA plate and incubated at 37°C for two days to allow metabolite accumulation, after which *P. e* was streaked into the adjacent quadrants and incubated for an additional day at 30°C. Two plates with *P. e* streaked alone served as controls. Plates were photographed and assessed for visible zones of growth inhibition to the *Enterococcus* streak. The assay was performed in two independent experiments with a total of12 plates.

#### Statistical analysis

All statistical analyses were performed in the statistical software R (Team, 2022). To assess microbiota population dynamics after host death, we fitted a zero-inflated generalized linear mixed-effects model with negative binomial error distribution, which were fitted using the package glmmTMB (Brooks et al., 2017), and model assumptions were assessed using the DHARMa package (Hartig, 2018). Microbiota load was the response variable, with square root transformed day post-death as fixed effect and replicate as a random effect. Pathogen load was analyzed using a linear mixed effects model with log-transformed pathogen load as the response. Fixed effects were day post death, the dietary treatment (CR and AB), and their interaction; replicate was included as random effect. To account for heterogeneity in error variances, we included diet and day as predictors of error variance using the option *dispformula* in the *glmmTMB* function.

The analysis of microbiota composition was performed using the R package *coda4microbiome* (Calle et al., 2023), which uses Compositional Data Analysis (CoDA) framework based on a log-ratio approach and penalized regression to identify taxa that increase and decrease in relative frequency. In our analysis we performed separate analyses for microbiota composition data obtained following snap freezing and death after *P. entomophila* injection. In each case we applied the function *coda_glmnet* for cross-sectional data, with day after death as a numeric covariate. The output of these analyses includes in each case a set of taxa that were identified to have increased or decreased in relative abundance during three days after host death. In addition, estimated coefficients provide information on how the magnitude of change in each taxon relates to the magnitude of changes in other selected taxa

#### Theoretical analysis

Day (2002) developed a mathematical model of pathogen virulence evolution that considers three transmission modes: 1) direct transmission from living hosts, 2) indirect transmission from living hosts via environmentally persistent spores, and 3) indirect transmission from dead hosts via spore release. Based on equation 4, Day derived a general expression for the conditions under which natural selection favors the evolution of increased pathogen-induced mortality: *γωDσ > β + κD σ*

where *γ* is the rate of pathogens clearance in living hosts, *ω* is the number of spores released at host death, *D* quantifies expected spore longevity in the environment (i.e. it is the reciprocal of the rate with which spores are degraded in the environment), *σ is* the transmission rate of environmental spores to susceptible hosts, *β* is the direct transmission rate between living hosts, and *κ* is the rate of spore release by living infected hosts.

This equation indicates that a higher transmission rate from dead hosts (*ω*) favors the evolution of pathogen-induced mortality. This transmission rate is expected to decrease when the host microbiota reduces pathogen proliferation after host death. Accordingly, competition between host microbiota and pathogens after host death should reduce selection for host killing and therefore act against a pathogen lifestyle of killing and exploiting dead hosts.

To illustrate this effect visually, we generated a Fig. 1 in the main text that is adapted from Fig. 1 in Day (2002). In this figure, Day illustrates the effect of variation in environmental spore longevity on the optimal pathogen replication *ε*, assuming that pathogen replication is the cause for infection-induced host mortality. Pathogen fitness *R* was given by (equation 3 of Day):

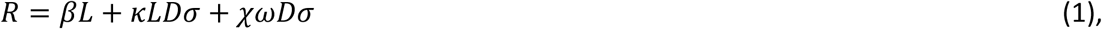

where *L* = 1/(*μ+ν+γ*) is the expected duration of an infection with *μ* being the host background mortality rate, and χ =(*μ+ν*)/(*μ+ν+γ*) is the probability that the infection ends due to host death instead of pathogen clearance. The three terms on the right-hand side in equation 1 correspond to direct transmission from living hosts, indirect transmission from living host, and indirect transmission from dead hosts.

For Fig. 1 Day assumed that virulence *ν*, direct transmission rate *β* and spore numbers after host death *ω* were functions of pathogen replication:

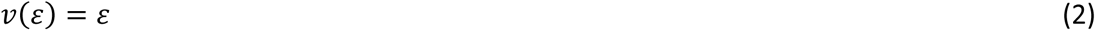

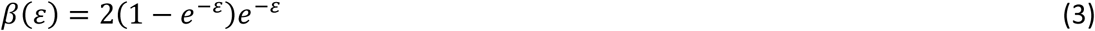

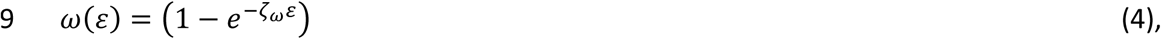

where *ζ_ω_*determines how spore release after host death and thus can capture how strongly the host microbiota limits spore release after host death. Additionally, we assumed that there is no spore release from living hosts (i.e. *κ*≡0), *σ*=0.1, *ζ_ω_*=5, μ=0.01 and γ=0.1. While Day explored two different values of *D* (0.2 and 10), we set D=10 and instead explored two different values of *ζ_ω_*: 1 and 10 (Fig. 4).

## Acknowledgements

We thank Jens Rolff and Sophie Armitage for their helpful comments on the manuscript, and Sophie Armitage for kindly providing the bacterial strain. We also thank Diana Aldana Alvarez, Luzie Mai Gallien, and Ylva Hüskes for their assistance with preliminary experiments, and Caroline Zanchi for helpful guidance regarding 16S rRNA sequencing. ChatGPT (OpenAI) was used intermittently to assist with text editing. Generated text was reviewed and revised by the authors and was not directly copied into the manuscript. This work was supported by the Deutsche Forschungsgemeinschaft (DFG), which provided funding to M.K. (KE 3013/1-1) and M.F. (FR 3061/6-1) as part of the Research Unit FOR 5026-2 InsectInfect. H.X. was supported by the China Scholarship Council.

## Author contributions

M.F. conceived the study; M.K. and M.F. designed the experiments; A.v.B. and H.X. performed the experiments; M.K., M.F., and A.v.B. analysed the data; M.F. performed the theoretical model analysis; and M.K. provided research guidance, and M.K. and M.F. wrote the paper.

