## supplementary material for "Microbiota-pathogen interactions after host death: a potential determinant of pathogen evolution"

### Supplementary materials

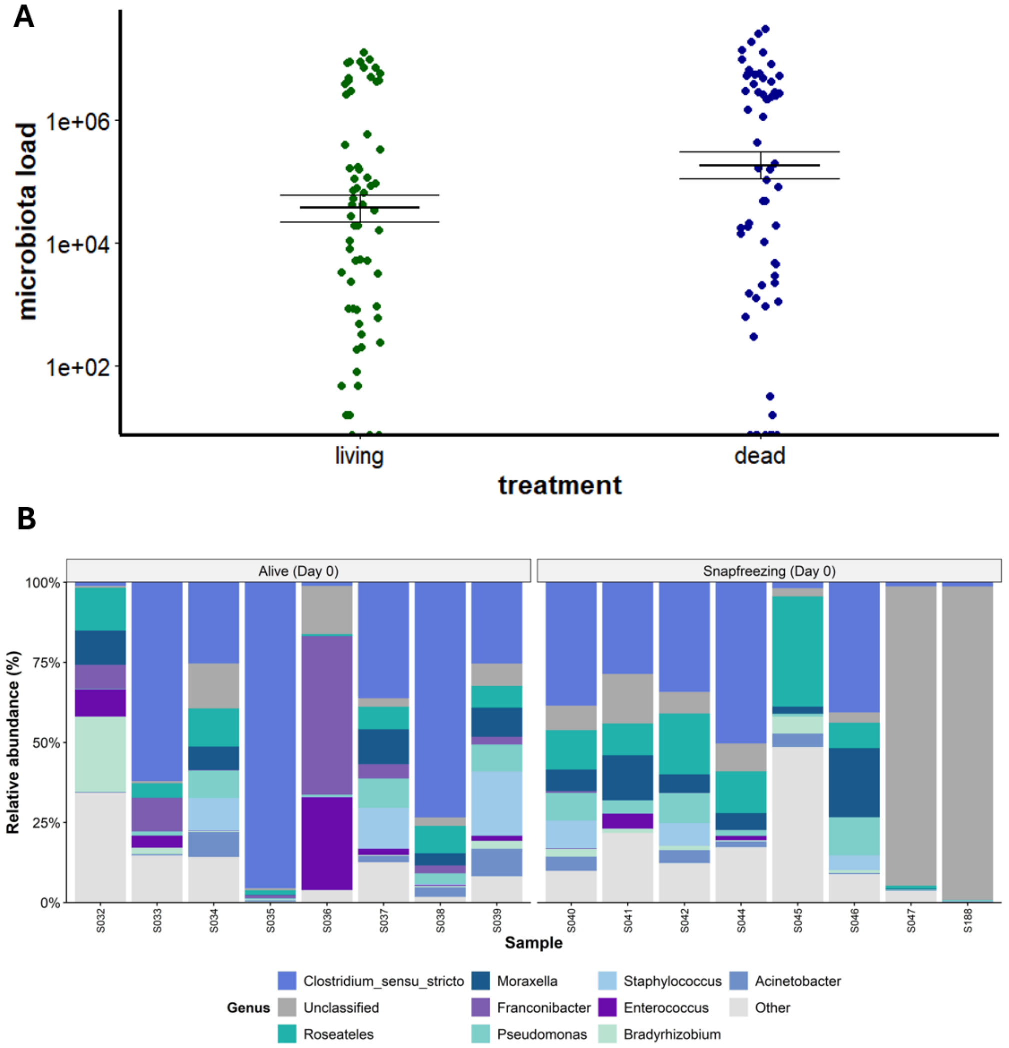


**Fig. S1:** Microbiota load in living larva and immediately after artificial death due to snap freezing. (**A**) Each dot represents the measured microbiota load of a single larvae, which were sampled in four independent replicates. Points were jittered on the x-axis to increase visibility. Bars indicate means and whiskers indicate corresponding standard errors. Results of a generalized linear mixed model revealed no indication of a statistically significant difference between microbiota load in living and snap frozen larvae (z=1.37, p=0.171). (**B**) Genus-level relative community composition of 10 dominant taxa in alive and snap-frozenlarvae at time zero, determined by 16S rRNA gene sequencing. Each bar represents an individual larva; colored segments denote distinct taxa, with segment height being proportional to relative abundance within that sample.

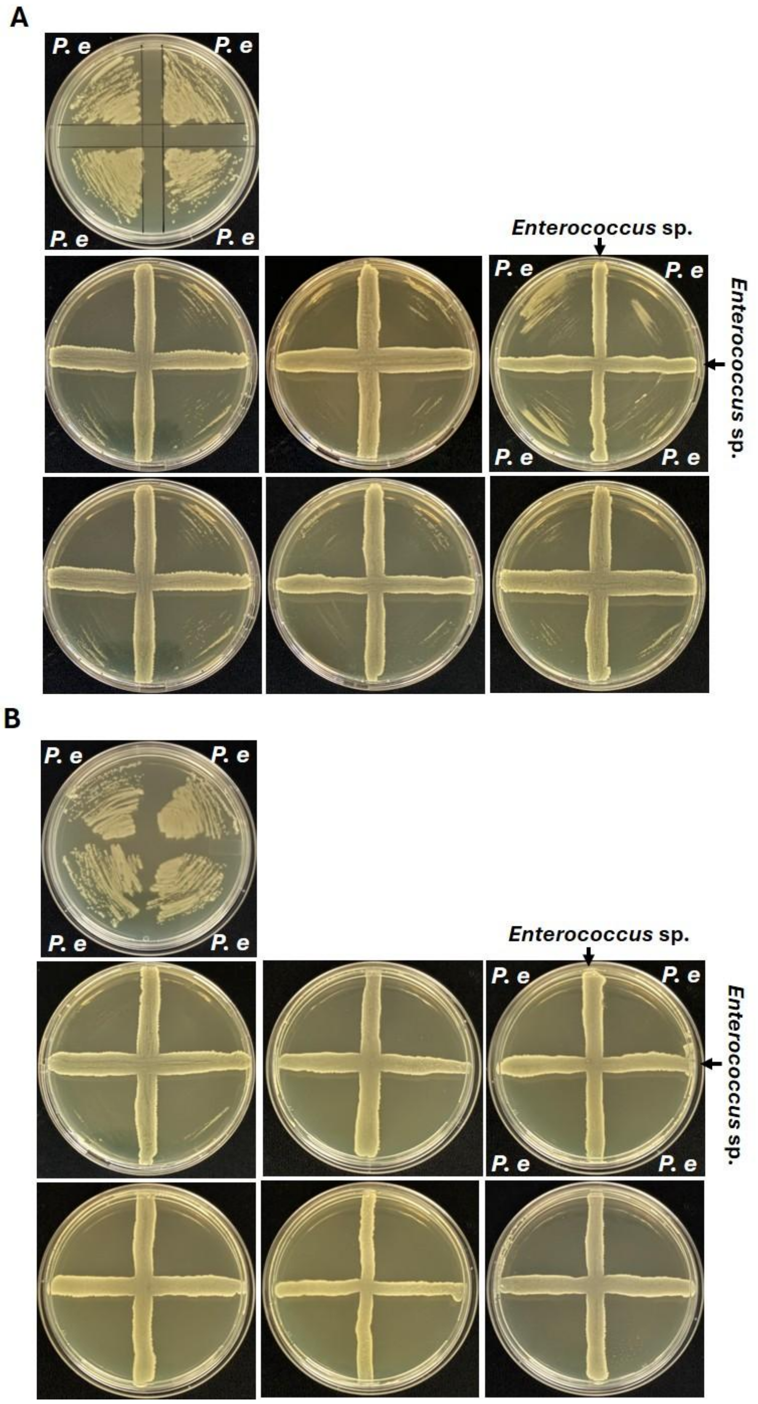


**Fig. S2:** Modified cross-streak assay of gut-derived *Enterococcus* sp. against *P. e.* (**A**) Biological replicate 1 and (**B**) replicate 2, representing inhibition of *P. e* growth by *Enterococcus* sp., indicated by a zone of inhibition adjacent to the *Enterococcus* streak. *P. e* grown in monoculture plates serves as control (top left in both panels).

**_Enterococcus_ sp. isolate S31 partial 16S rRNA gene sequence**

CCGCGGTAATACGTAGGTGGCAAGCGTTGTCCGGATTTATTGGGCGTAAAGCGAGCGCAGGCGGTTTCTTAAGTCTGATGTGAAAGCCCCCGGCTCAACCGGGGAGGGTCATTGGAAACTGGGAGACTTGAGTGCAGAAGAGGAGAGTGGAATTCCATGTGTAGCGGTGAAATGCGTAGATATATGGAGGAACACCAGTGGCGAAGGCGGCTCTCTGGTCTGTAACTGACGCTGAGGCTCGAAAGCGTGGGGAGCAAACAGG
